# A recombinant CHIKV-NLuc virus identifies chondrocytes as target of Chikungunya virus in a immunocompetent mouse model

**DOI:** 10.1101/2024.05.20.594924

**Authors:** Vincent Legros, Essia Belarbi, Patricia Jeannin, Beate Kümmerer, David Hardy, Philippe Desprès, Antoine Gessain, Pierre Roques, Pierre-Emmanuel Ceccaldi, Valérie Choumet

## Abstract

First isolated in 1953 in Tanzania, the arthritogenic Chikungunya virus (CHIKV) re-emerged globally in 2005, leading to widespread outbreaks. Unlike other arboviruses, CHIKV predominantly induces symptomatic infections (72-96%), marked by fever, myalgia, polyarthralgia, and rash. Although rarely fatal, atypical forms such as encephalopathies can occur. Notably, 75.4% of patients experience persistent arthralgias for up to three years after the acute phase. Understanding CHIKV’s pathophysiology in the joints is challenging due to the difficulties to obtain biological samples. The study employs a mouse model infected with a reporter virus expressing a Nano Luciferase to investigate the disease’s transition to chronic arthritis. The murine model reveals viral replication in metatarsi joints, particularly in chondrocytes, confirmed in primary human chondrocytes undergoing viral-induced apoptosis. *Ex vivo* analysis confirmed viral replication in leg bones and articular cartilages, with histological evidence of focal erosive lesions and periarticular inflammation. The study further utilizes an *in vivo* imaging mouse model to monitor viral replication over time. Human chondrocytes prove susceptible to CHIKV infection, exhibiting active viral replication, bioluminescence activity, and increased viral production. CHIKV induced apoptosis, the upregulation of markers associated with cartilage remodeling and altered the cytokine production. This comprehensive study, utilizing advanced techniques and models, provides insights into CHIKV’s ability to infect articular cartilages, shedding light on the mechanisms of chronic arthritis following infection.

## Introduction

Chikungunya virus (CHIKV) is an arthritogenic arbovirus belonging to the *Togaviridae* family, *Alphavirus* genus and responsible of incapacitating acute and chronic musculoskeletal disease [1]. Chikungunya virus genome is a single stranded positive RNA virus of approximately 11.8Kb, with two open reading frames (ORFs) encoding non-structural proteins (nsP 1–4) and five structural proteins, among which the capsid protein and two envelope glycoproteins E1 and E2 [2]. The virus was first isolated in 1953 from a patient having a symptomatic picture of fever and arthralgia in Tanzania [1]. CHIKV has re-emerged in 2005 in the Indian ocean and was responsible since of several outbreaks around the world involving millions of human cases [3].

CHIKV infection is particularly notable because, unlike other arboviruses, infected individuals are predominantly symptomatic (72-96%) [4]. Symptoms occur after an average incubation period of three days and usually consist in an abrupt onset of fever, followed by myalgia, polyarthralgia and macupapular rash. The arthralgia is often symmetrical and can affect any joint. Synovitis has been reported in 32–95% of patients [5]. Although CHIKV infection is rarely fatal, atypical forms of the disease (e.g. encephalopathies, myocarditis, hepatitis, and multi-organ failure) have been observed [6]. Viral RNA is detectable in the blood at the onset of symptoms and can be found in very high quantities (>10^9^ copies per milliliter), allowing for new human-to-mosquito transmission cycles [7]. Viremia is usually controlled after the seventh day of infection, concomitant with the appearance of anti-CHIKV immunoglobulins in the blood [8].

One of the most remarkable features of CHIKV infection is the high proportion of patients presenting persistent manifestations of the disease after the resolution of the acute phase (7-10 days) [9]. Chronicity is characterized by muscle and joint pain located most frequently in the wrists, ankles, knees, metacarpal and metatarsal joints lasting up to 3 years [10]. Many epidemiological studies have attempted to quantify the percentage of patients with a chronic form of the disease. As an example, a study conducted after the emergence of CHIKV in Reunion Island reported that 75.4% of patients suffered from persistent arthralgias after a median duration of 1.5 years [11]. This persistent arthralgia is often debilitating but in most cases, gradually improves until complete resolution. Several hypotheses may be put forward concerning the pathogenesis of these manifestations: viral persistence in the joint with cytopathic effect due to replication, persistence of viral antigens, immune-mediated response, or development of an autoimmune response after the infection [12].

The pathophysiology of CHIKV infection remains poorly understood, despite numerous studies. One of the main reasons is the extreme scarcity of available biological samples. As a matter of fact, while human blood samples are commonly available, synovial fluid collection is an invasive procedure and therefore samples are very rare. Furthermore, for obvious ethical reasons, rare studies analysed biopsies of patients’ joint tissue [13,14]. In the absence of joint samples, the use of animal models is the only method to understand a phenomenon as complex as alphaviral arthritis.

Several animal models of CHIKV infection have been developed [8][15–20] making it possible to characterize the infection and associated histopathological lesions of the spleen, liver, lymph nodes and muscle, and the development of peri-articular oedema in the acute phase of infection. Some models also allow the study of CHIKV chronic infection [18,21–24] as it has been suggested to occur in humans [13,14,25]. A study on partially immunodeficient mice [16] identified fibroblasts of the dermis, muscle and joint capsule, muscle satellite cells, epithelial and endothelial cells of many organs such as liver, brain and spleen as CHIKV cellular targets during the acute phase. In a simian model, liver endothelial cells and macrophages have been found immunoreactive for CHIKV antigens during the chronic stage [18].

Little is however known about the cellular targets of CHIKV in humans. *In vitro*, human cell lines such as epithelial, fibroblastic and some endothelial cells were found susceptible to CHIKV infection as well as primary cells such as macrophages, keratinocytes, synovial and dermal fibroblasts, ostoblasts, chondrocytes, myoblasts [26–28]. Very scarce *in vivo* data are available, except the demonstration of fibroblast infection in the dermis, joint capsule and muscle facia from a single fatal neonatal case [16], and satellite muscle cells in biopsies from two adult patients with myositis [28].

Very few studies have focused on cellular targets and infection characteristics in the joint. In murine models, only two studies showed the presence of viral antigens and/or viral RNAs in the joint during the acute phase, and viral RNA, without infectious particles, during the chronic phase [22,24,29]. In infected patients with chronic arthrtitis, viral antigens could be detected in cells from joint tissue, without cell type characterization [13], and in synovial macrophages [14]. However, the detection of viral RNAs or antigens from the joint does not answer several major questions. The first concerns the tropism of the virus. Indeed, the ability of this virus to remain in the joint implies that it is capable of setting up a persistent infection or periodically reinfect new cells, while limiting the immune response.

The objective of this study was to better characterize the mechanisms of transition to chronic arthritis related to CHIKV infection, using a previously published mouse model [15] and a bioluminescent reporter virus. Our hypothesis was that the phenomenon of viral persistence takes place within a specific tissue, the articular cartilage. In order to explore this, we used a new tool that was generated in our laboratory, a reporter virus expressing Nano Luciferase (NLuc) which allowed us to successfully develop an *in vivo* imaging mouse model of CHIKV acute and chronic infection. Our strategy in using these techniques was to precisely localize the sites of replication and persistence of the virus, in an immunocompetent mouse model, in combination with more classical techniques such as histology. Moreover, the use of primary human cells allowed to confirm the results obtained *in vivo* in order to extend them to humans, as well as to characterize more precisely the mechanisms involved. By this approach, we showed here that metatarsi joints harbored viral replication after the end of the acute phase. We identified chondrocytes as the major targets of infection, and observed viral-induced apoptosis in primary human chondrocytes.

## Material and Methods

### Design of the recombinant CHIKV-NLuc

A Nano Luciferase reporter gene was inserted in-frame within the Spe I restriction site of a CHIKV infectious clone, based on a CHIKV isolated in Mauritius in 2006 (BNI-CHIKV 899; accession number: FJ959103.1) [30]. For this purpose, a previously described strategy [31,32] was used; briefly, the NLuc reporter gene was inserted in-frame between nsP3 and nsP4 in the non-structural polyprotein precursor and flanked on both sides by nsP2 protease cleavage sites. Viral RNA was *in vitro* transcribed from the linearized recombinant infectious clone using the mMESSAGE mMACHINE Kit (Ambion) and electroporated into Vero E6 cells. Viral supernatants were collected 48 hours post inoculation (hpi) and viral stocks for experiments were produced after a second passage. All viral stocks produced were titrated by plaque assay on Vero E6 cells. The resulting virus was named CHIKV-NLuc in which the reporter is expressed as part of the non-structural polyprotein precursor and is cleaved by the viral protease nsP2 during the replication cycle (GMO agreement 2454 from the French Ministry MESRI). In some experiments, the wild type CHIKV strain (CHIK 05-049, accession number: AM258994.1) was used.

### NLuc activity quantification

Cells or tissues were lysed with Passive Lysis Buffer (Promega, USA) before centrifugation for 10 minutes at 13,000g followed by a ten times dilution in Phosphate-buffered saline (PBS). Each sample was mixed volume to volume with a 1% furimazine solution. NLuc activity was measured with a Centro LB 960 luminometer (Berthold Technologies, Germany) and expressed as relative luminescence unit (RLU).

### Assessment of CHIKV-NLuc stability

To assess the stability of the recombinant virus, Vero E6 cells were seeded and infected with CHIKV-NLuc at a multiplicity of infection (MOI) of 1. Twenty-four hours after infection, 100µL of supernatant was used to infect new cells. The remainder of the supernatant was retained to assess the infectious titre. The cells were washed once with PBS and quantification of NLuc activity was performed on the cell lysates. This operation was repeated for a total of 10 successive passages.

### Cells

Vero E6 cells (ATCC CRL-1586) were used for viral production and titration. Human chondrocytes (HC) (Ref PB-402-05a) were cultured according to supplier’s instructions (Cell Applications, USA). The cells were incubated at 37°C and 5% CO_2_.

### Cell infection

For *in vitro* infections, confuluent cell monolayers were seeded in the appropriate culture medium. Aliquots of virus were diluted to obtain the targeted MOI (0.1, 1 and 10) for inoculation. After an hour, the inoculum was removed and the cells washed once before incubation until collection.

### Viral titration

Plaque assay titrations were performed on Vero E6 cells. Briefly, confluent monolayers of Vero E6 cells were seeded and inoculated with 1:10 serial dilutions of the sample. After an hour of incubation, unadsorbed virus was removed and DMEM medium supplemented with 1.6% carboxymethyl cellulose (CMC) and 2% Fetal Calf Serum (FCS) was added. The supernatant was removed after 2 days and the cells were washed and fixed with a crystal violet solution (PBS with 4% paraformaldehyde and 0.2% crystal violet). The fixed cells were finally washed with water and the number of lysis foci was counted to assess the viral titre. The results were expressed as plate-forming units per mL (pfu/mL).

### Mice

Animal experiments were approved by the Ethics Committee of the Institut Pasteur and authorised by the French Ministry of Higher Education and Research (reference CETEA 0762.02). The animal manipulations involving wild type strains of CHIKV were performed on four-week-old female C57Bl/6 mice (Charles River, France). In order to limit the absorption of the bioluminescent signal due to the black colour of the coat, four-week-old albino female C57Bl/6N (B6N-TyrC-Brd/BrdCrCrl) mice (Charles River, France) were used for the experiments involving CHIKV-NLuc.

Mice were anesthetized with an intraperitoneal ketamine and xylazine solution and inoculated subcutaneously with CHIKV or vehicle into the footpad of the right paw. The volume injected was 25µL for an infectious dose of 10^3^, 10^4^, 10^5^ or 10^6^ pfu. After infection, the mice were observed daily. Weight, clinical signs, plantar oedema, viremia and blood parameters were assessed.

Blood samples were obtained by puncture at the distal end of the tail, for a volume strictly inferior to 30µL. Larger blood volumes were taken by intracardiac punctures during terminal sampling and under deep anaesthesia. Blood parameters were monitored using the Vet ABC™ Hematology Analyzer (Scil, Germany). To perform tissue sampling, mice were anaesthetised and sacrificed by intra-cardiac infusion of PBS. Tissue samples intended for primary cell isolation or bioluminescent signal quantification were placed in PBS and kept at 4°C. Soft tissues (skin and muscle) were removed by dissection to expose bones and joints. The cartilages of the hip, knee and metatarsals were then isolated by dissection under a binocular loupe. For *ex vivo* bioluminescent signal reading, the joints were placed in passive lysis buffer (Promega, USA) and mechanically ground with ceramic beads (Precellys, Bertin Technologies, France).

### Histopathology

After sacrifice, legs were removed, fixed for 48 to 72 hours in 10% neutral buffered formalin, and embedded in paraffin; 4-μm-thick sections were performed and stained in hematoxylin-eosin or safranin. Sections were analyzed by a trained veterinary pathologist by separate blinded scoring.

### Viral replication assessment

Total RNA was extracted using the Nucleospin® RNA II kit (Macherey-Nagel, Germany) according to the manufacturer’s instructions, eluted in 70µL of RNAse-free water and stored at -80°C before analysis. Relative quantification was used for cellular RNAs using the GAPDH housekeeping gene as reference. For viral RNA, copy numbers were assessed with a quantitative reverse transcription polymerase chain reaction (RT-qPCR) using the Power Sybr Green RNA-to-Ct one step kit (Applied Biosystems), a dilution of synthetic CHIKV RNA as standard, and the following primers described in a previous publication [33]:

CHIKV/E2/9018 sense primer (CACCGCCGCAACTACCG)

CHIKV/E2/9235 anti-sense primer (GATTGGTGACCGCGGCA).

The signal was normalised relative to the standard curve. For ΔCt analysis, the normalised data were used to estimate the RNA copy number in each well.

### Immunofluorescence

Cells were grown on coverslips, infected with different CHIKV MOIs (0.1, 1, 10), fixed at different times with a 4% paraformaldehyde solution and a permeabilized with 0.1% Triton X-100 solution. The cells were washed and incubated with the 3E4 recombinant anti-CHIKV E2 murine antibody (Creative Biolabs, USA). When indicated, an anti-collagen type II rabbit antibody (Novus Biologicals, USA) was used. Anti-mouse IgG and anti-rabbit IgG coupled respectively with Cy5 and AlexaFluor 488 dyes, were used as secondary antibodies. The coverslips were mounted with ProLong gold antifade reagent with DAPI (Life Technologies, USA) and were examined using a fluorescence microscope (EVOS, Thermofischer Scientific, USA).

### TUNEL apoptotic and caspase 3/7 activity assay

Human chondrocytes were infected with CHIK MOI 1 and 10. Cells were processed at different times post inoculation (pi) according to manufacturer’s instructions (Caspase-Glo® 3/7 Assay kit, Promega, USA) to determine caspase 3/7 activity. Luminescence of each sample was analyzed with a Centro LB 960 luminometer (Berthold Technologies, Germany). The caspases activity was expressed as RLU.

A terminal deoxynucleotidyl transferase (TdT) dUTP nick-end labelling (TUNEL) assay was performed using the *In Situ* Cell Death Detection Kit, Fluorescein (Roche, Switzerland) according to manufacturer’s instructions. Briefly, HC were seeded on coverslips and infected with CHIKV (MOIs of 1 and 10). The cells were subsequently fixed at 24 and 48 hpi with 4% paraformaldehyde solution and endogenous peroxidase activity was blocked using a 3% H_2_O_2_ solution in methanol. Cells were then permeabilised with a 0.1% Triton-X and 0.1% sodium citrate solution. Negative labelling controls (without TdT) were included, along with mock-infected controls. Samples were subjected to labeling with the TUNEL reaction mixture prior to imaging analysis. Coverslips were mounted with ProLong gold antifade reagent with DAPI, and examined using a fluorescence microscope (EVOS Thermofischer Scientific, USA).

### Cytokine relative quantification in the supernatant

Supernatants from infected and uninfected primary cells were collected and centrifuged for 10 minutes at 13,000g to remove cell debris. The relative levels of cytokines were assessed using the Proteome Profiler Human XLCytokine Array kit (R&D Systems, USA) according to the manufacturer’s instructions. Membranes were analysed using myECLImager and bioluminescent signals quantified with myImageAnalysis version 1.1 (Thermofischer Scientific, USA). Signal from infected cells’ supernatants were normalised to negative controls and expressed as percentage change.

### In vivo imaging

Mice inoculated with virus or diluent alone were anaesthetised and then injected with 4mg/kg of furimazine. *In vivo* imaging was performed using the IVIS spectrum system and the data obtained were analysed using Living Image 4.5 software (Perkin Elmer, USA). In order to quantify bioluminescence, regions of interest (ROI) of identical area and shape were manually defined. The results were then expressed as total flux (photons per second per square centimetre per steradian: photons/s/cm^2^/sr).

### Isolation of primary murine chondrocytes

The cartilages isolated under binocular microscope were dilacerated using a sterile scalpel, then incubated in a 0.5mg/mL trypsin solution (Sigma-Aldrich, USA) under agitation at 37°C for 30 minutes. Following centrifugation, the solution was removed and replaced with 0.1% collagenase II in DMEM with 5% FCS and samples were incubated for 4 hours at 37°C with agitation. The samples were then placed on 40µm filters (Cellstrainer, Thermo Fischer Scientific, USA) and centrifuged at 500g for 5 minutes. The isolated cells were resuspended in DMEM supplemented with 5% FCS and immediately plated on glass coverslips using Cytofunnels (Thermo Fisher Scientific, USA) and centrifugation at 1500g for 10 minutes with a Cytospin 4 (Thermo Fisher Scientific, USA). The plated cells were fixed with a 4% paraformaldehyde solution and labelled for immunofluorescence with the anti-CHIKV 3E4 and anti-collagen type II antibodies.

### Statistical analyses

Statistical analyses were performed using Prism version 6 software. Spearman’s correlation test was used to investigate relationships between variables. Non-parametric Mann-Withney tests and ANOVA followed by Bonferroni correction were used to test for differences between groups. P-values less than 0.05 were considered as significant.

## Results

### Assessment of CHIKV-NLuc stability

We first explored the stability of the viral construction (Fig. 1A). Conservation of the transgene NLuc was tested by serial infections and assessment of the viral titer and luciferase activity. As shown in Fig.1B, the bioluminescent signal is remarkably stable for 10 serial passages, with however a one log decrase in the luciferase activity at the last passage. Nevertheless, we observe a strong positive correlation between the bioluminescence and the viral titer (Spearman correlation test, r=0.93, p-value=0.0007). CHIKV-NLuc appears to be stable and can be used to monitor the infection by *in vivo* imaging.

**Figure 1.**
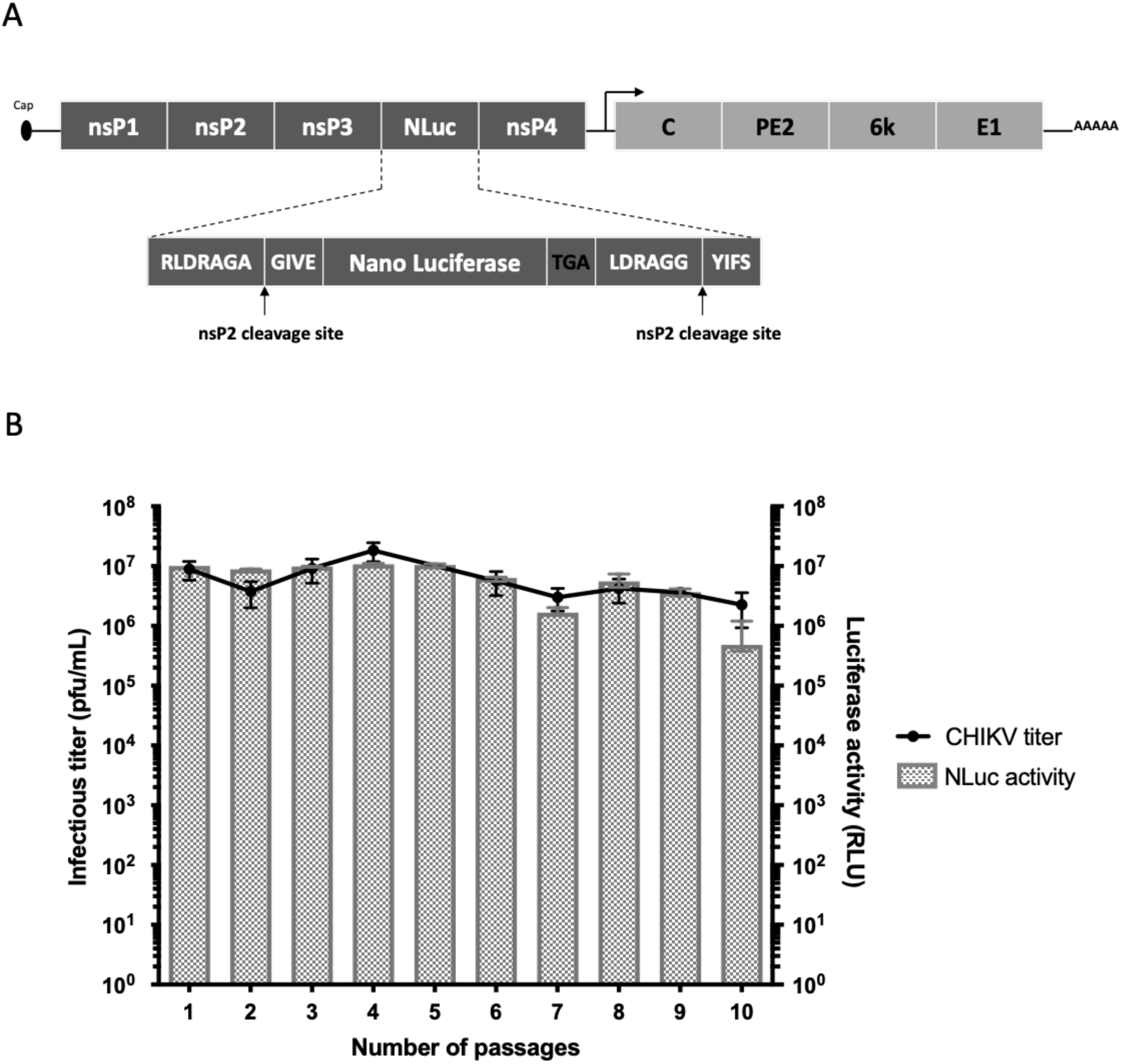
Generation of a recombinant CHIKV expressing NLuc. (A) Genome organization of the recombinant CHIK-NLuc virus. NLuc gene was inserted in-frame between nsP3 and nsp4 genes. Arrows represent nsP2 clivage sites. (B) Stability of the recombinant CHIK-NLuc virus. Vero E6 cells were serially infected for 10 passages with CHIKV-NLuc to assess the stability of the construct. At each passage, the viral titer of the supernatant was measured (blue curve), as well as the luciferase activity (red bars), expressed as RLU in the infected cells. The curves and histograms show the median value and interquartile range of measurements from 4 replicates.

### CHIKV-NLuc allows the visualization of viral replication sites during infection in an immunocompetent mouse model

We adapted the immunocompetent mouse model of infection previously described by Gardner et al. [15] using the recombinant CHIKV-NLuc virus. Albino C57Bl/6 mice were subcutaneously infected in the footpad, and viral replication was assessed at different times post-inoculation (pi) by bioluminescence assay after injection of the NLuc substrate (Fig. 2).

**Figure 2.**
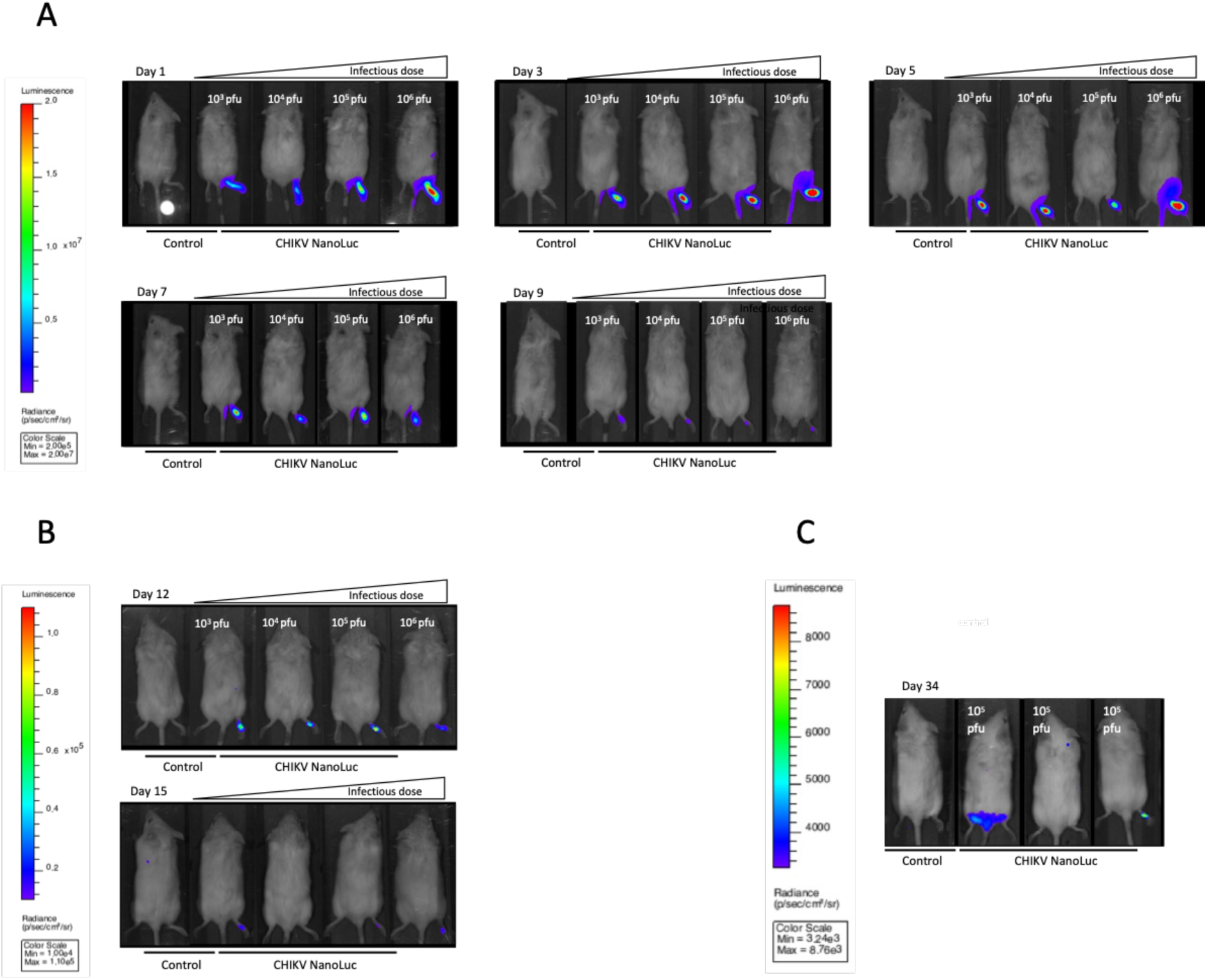
Plantar pad infections with CHIKV-NLuc generate sufficient bioluminescent signal for monitoring. Mice were inoculated into the plantar pad of the right hind paw with the diluent alone (control, on the left) or CHIKV-NLuc (10^3^ to 10^6^ pfu/mouse, from left to right). The bioluminescent signal intensity is associated with a colour code according to the scale indicated at the right of the images. (A) Mice at 1,3,5,7 and 9 dpi (with a scale of 10^6^ to 2.10^7^ p/sec/cm^2^/sr). (B) Mice at 12 and 15 dpi (scale of 10^4^ to 10^5^ p/sec/cm^2^/sr). (C) Mice at 34 dpi (scale of 10^3^ to 10^4^ p/sec/cm^2^/sr).

As shown in Fig. 2A, a bioluminescent signal is detected in the inoculated footpad, as early as the 1^st^ day after inoculation. During the acute phase, the signal increased until 5 days pi (dpi), extended into the paw until the hip joint and gradually decreased at 7 to 9 dpi. A bioluminescent signal was also detected in the rest of the body, however it is not displayed on the images as the very high signal from the inoculated paw would saturate the camera. During the post acute phase (12 and 15 dpi, Fig. 2B), a weak signal persisted in the inoculated footpad. Interestingly, during the chronic phase, at 34 dpi (Fig. 2C), a bioluminescent signal could still be observed in two mice.

We thus performed quantification of the bioluminescent signal for the whole body, the inoculated paw and the contralateral paw. In the inoculated paw as well as for the whole body, the signal is significantly higher until 7 dpi compared to uninfected mice (Fig. 3 A, B). For the inoculated paw the signal remains higher than uninfected controls at 9 dpi, although not statistically significant. In the contralateral paw (Fig. 3C) the signal is significantly increased only at 5 dpi as observed for whole body or inoculated paw, and decreased at 9 dpi. At 30 dpi no significant signal could be detected using this technique (A, B, C).

**Figure 3.**
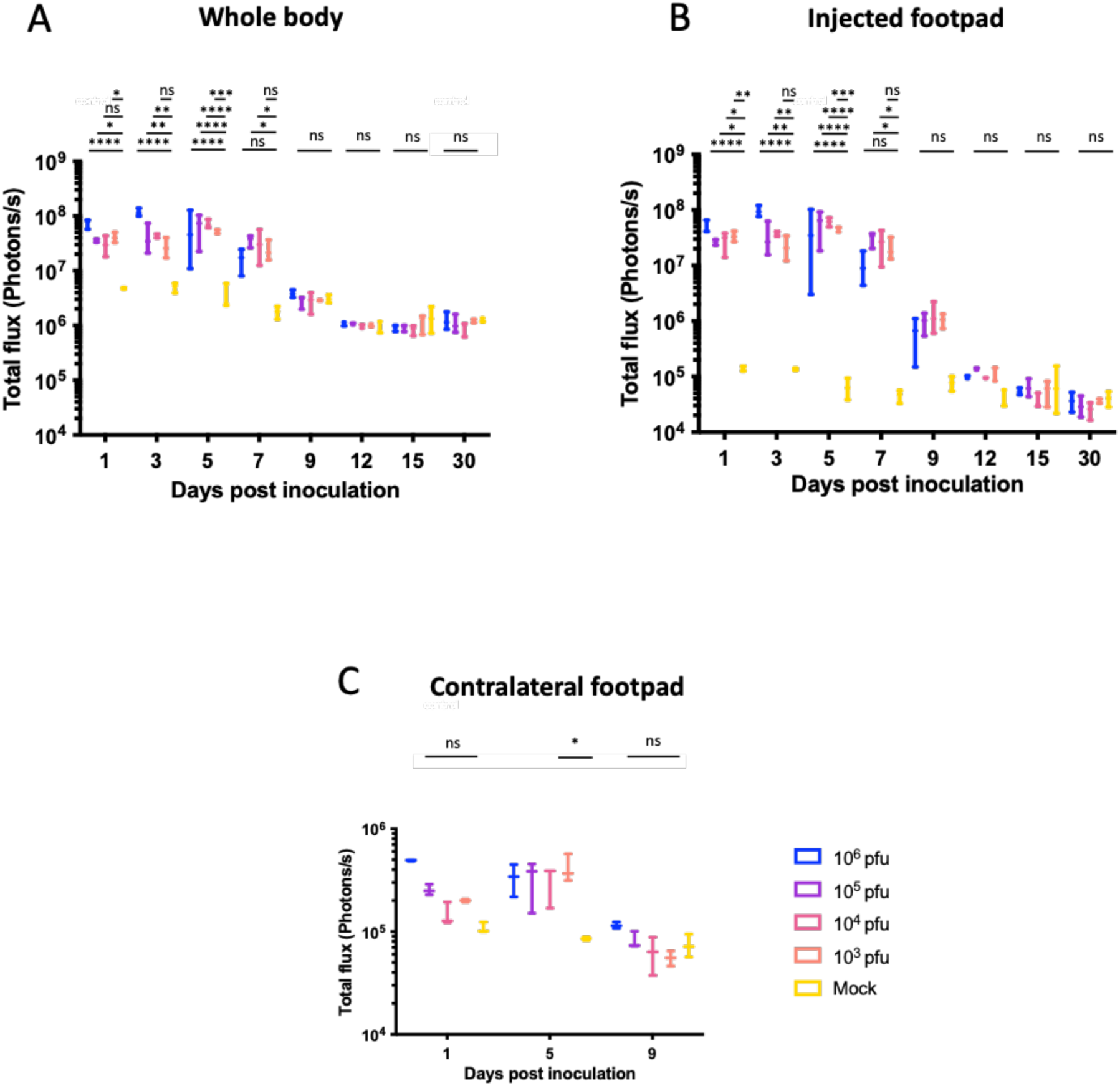
Assessment of bioluminescent signal in CHIKV-Nluc infected mice. (A) whole body, (B) inoculated paw and (C) contralateral paw of mice inoculated with 10^6^ pfu (blue), 10^5^ pfu (red) 10^4^ pfu (green), 10^3^ pfu (purple) of CHIKV-NLuc or non-infected (orange). Mice were imaged, regions of interest of the same area and location were defined and the signal quantified at 1, 3, 5, 9, 12, 15 and 34 dpi and expressed as total flux in photons per second. Median and interquartile range of measurements from three mice are given. ns: not significant, *: p-value < 0.05; **: p-value < 0.01; ***: p-value < 0.001; ****: p-value < 0.0001 (Analysis in double variance).

The bioluminescent signal observed in Fig. 2 indicated viral replication in the ankle/metatarsus joints. However, the resolution was not sufficient to allow an accurate identification of the tissues involved in such replication, especially in later stages (30 dpi). In order to increase the specificity of the measure, *ex vivo* signal quantifications were performed. Mice were infected as previously described. First, leg bones of an uninfected and infected animals (12 dpi) were isolated. Bioluminescence imaging revealed a significant signal in the leg of the infected animal (Fig. 4A). In order to perform a quantification, mice were euthanized at 6, 12 and 30 pi and perfused with PBS to remove blood from tissues. Using a binocular magnifier, articular cartilages from hips, knees and ankles/metatarsis were dissected and grinded. The NLuc activity was quantified and expressed as fold change relative to uninfected samples. As shown in Fig. 4B, the luciferase activity was significantly higher at 6 dpi in the cartilages of infected mice, indicating viral replication in the articular cartilage. Indeed, we observed an increase in luciferase activity up to 10^3^ fold in ankle/metatarsis cartilages. Interestingly, while the knee and hip cartilages from infected mice shows no difference compared to the control at 12 and 30 dpi, those from the ankle displayed a significantly higher signal compared to uninfected cartilages. Legs from CHIKV-infected mice collected on 6, 12 and 30 dpi were fixed in formalin and embedded in paraffin. At 6 dpi, after safranin staining, in comparison with the cartilage of control animals, (Fig. Suppl1 A, B), focal erosive lesions were observed in the cartilage of the metatarsophalangeal joints, indicating limited destruction of this tissue, (Fig. Suppl. 1 C, D). Interestingly, using 19emalum eosin staining and in contrast to uninfected control (Fig. Suppl1 E), an inflammatory infiltrate of the periarticular connective tissue and the joint capsule was visible at 6 dpi in the metatarsophalangeal joint (Fig. Suppl1 F). At 12 dpi, persistent periarticular inflammation was visible, without marked destruction or regeneration of articular cartilage (Fig. Suppl1 G). At 30 dpi, a mild periarticular inflammation was still visible (Fig. Suppl1 H).

**Figure 4.**
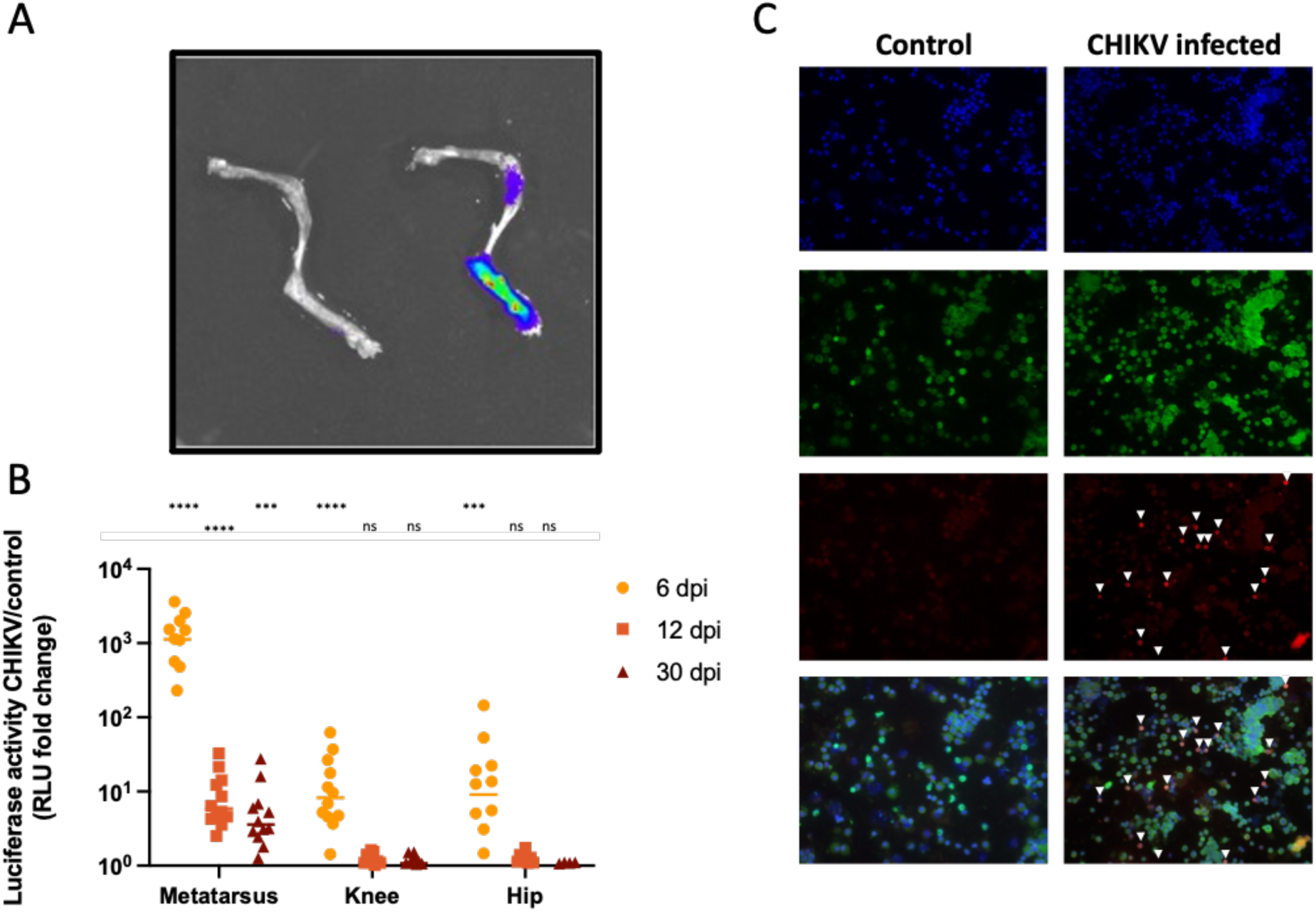
*Ex-vivo* visualization and measurement of luciferase activity and infection in the leg bone and the articular cartilage of the ankle/metatarsals, knees and hips of infected mice. Mice were inoculated into the footpad of the right hind leg with CHIKV-NLuc (10^5^ pfu) or the diluent alone (control). (A) Leg bones of control and infected animals were dissected at 12 dpi, and analyzed by bioluminescence imaging. Left: uninfected; Right: infected. (B) Using a binocular magnifying glass, articular cartilages (hip, knee, ankle/metatarsis were isolated at 6, 12 and 30 dpi and luciferase activity was quantified. Each point represents the luciferase activity of a cartilage normalized to uninfected controls and measured in two independent experiments, median and interquartile range are also shown. Ns: not significant, *: p-value < 0.05; **: p-value < 0.01; ***: p-value < 0.001; ****: p-value < 0.0001 (non-parametric Mann-Withney test). (C) Primary murine cells isolated from metatarsal cartilage at 6 dpi. The arrows indicate cells with double labelling of type II collagen and viral E2 protein.

### *In vivo* infection of murine chondrocytes

The *in vivo* bioluminescence imaging suggested active replication of CHIKV-NLuc in the articular cartilages. In order to increase the specificity of our analysis and to characterize the infected cell type, infected mice were euthanized at 6 or 30 dpi, and their legs were dissected. Ankle/metatarsus and knee cartilages were isolated as previously described and dissociated using trypsin and collagenase II treatments. To avoid any replication resulting from infection after isolation, the primary cells were immediately plated and fixed on a slide with a solution of 4% paraformaldehyde. CHIKV E2 protein and type II collagen, a chondrocytes marker, immunoreactivities were assessed. Strikingly, at 6 dpi, most of cells from ankle cartilages showed immunoreactivity for the CHIKV E2 protein and type II collagen (Fig. 4C). In contrast, only few type II collagen positive cells were found to be also immunoreactive for E2 in the knee (data not shown).

Interestingly, a limited number of infected chondrocytes (E2 and collagen II immunoreactive) could still be detected as late as 30 dpi in the metatarsal articular cartilage (Fig. 5).

**Figure 5.**
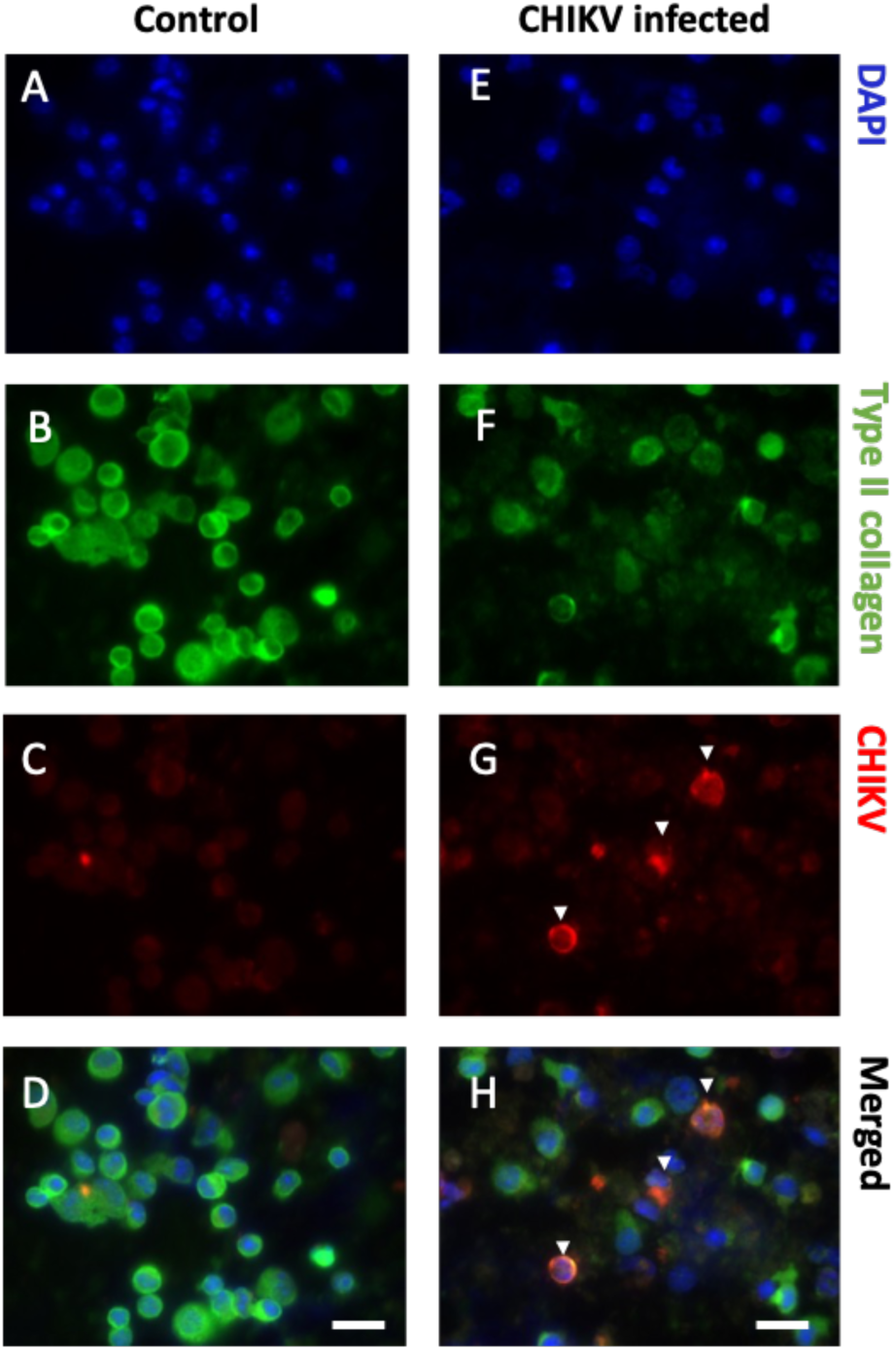
*Ex-vivo* visualization of infection in metatarsal articular cartilage of infected mice at 30 dpi. Mice were inoculated into the footpad of the right hind leg with CHIKV-Nluc (10^5^ pfu) or the diluent alone (control). Primary murine cells isolated from metatarsal cartilage at 6 dpi. The arrows indicate cells with double labelling of type II collagen and viral E2 protein. D, F: merge of Dapi, collagen II and E2 staining: Scale Bar: 20µm.

### Human chondrocytes are susceptible to CHIKV infection

We then investigated the susceptibility of primary human chondrocytes (pHC) to CHIKV. In order to evaluate the viral replication in pHC, we performed *in vitro* infection with CHIKV 05-049 strain at the MOIs of 0.1, 1 and 10 or CHIKV-NLuc, MOI 1. Viral RNA was detected in infected cells as early as 4 hpi. (Fig. 6A). At 8 hpi the viral RNA quantity was 2 logs higher than at 4 hpi, before reaching a plateau between 12 and 24 hpi at 10^8^ RNA copies/5.10^4^ cells (Fig. 6A), indicating active replication. Bioluminescence activity was observed in CHIKV NLuc infected pHC following a similar kinetics as wild type strain (Fig. 6B). Moreover, we confirmed using plaque assay titration that the infection of pHC led to the production of infectious viral particles, with a peak at 36 hpi (10^7^ pfu/mL) (Fig. 6C). CHIKV infection was also assessed using an immunofluorescence assay: immunoreactivity for the viral glycoprotein E2 could be detected as soon as 1 dpi. (Fig. 6D).

**Figure 6.**
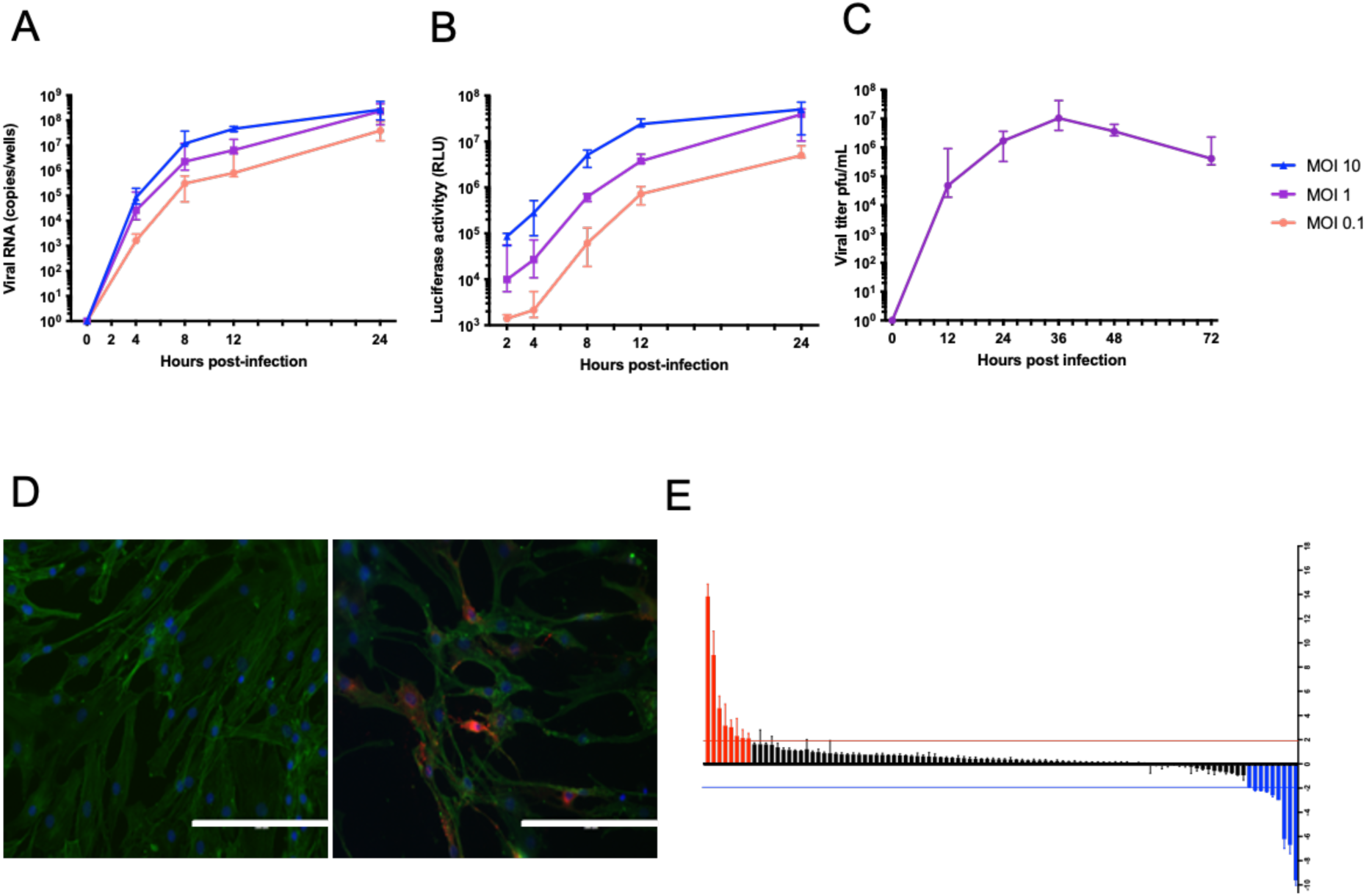
Infection of primary human chondrocytes with CHIKV. (A) Assessment of viral RNA levels in CHIKV infected pHC. Viral RNA was quantified in the cells lysates by RT-qPCR at different times pi and MOIs; (B) Bioluminescence activity, expressed in RLU, after infection with CHIKV NLuc, at three different MOI; (C) Number of infectious virus particles in the supernatant of CHIKV-infected pHCs assessed by plaque assay. The curves show the median value and interquartile range obtained from three independent experiments. (D) In vitro cultured pHCs were inoculated with CHIKV (MOI 1) or vehicleobserved by fluorescence microscopy. (Scale Bar: 200 μm). Left: uninfected cells; right: CHIKV infected cells. E: Effect of CHIKV infection of pHCs on cytokine production. Supernatants from CHIKV-infected and uninfected pHCs were analysed at 6 hpi. Threshold corresponding to a 2 fold increase or 2 fold decrease compared to control cells ( highlighted red and blue bars respectively).

### CHIKV infection of pHC induces apoptosis and early modifications in cytokine production

To explore the effect of CHIKV on the physiology of chondrocytes, we monitored the change in cytokine production following infection. Cytokines present in the supernatant were investigated using a broad-spectrum immunoenzymatic assay at 6 hpi. On the 102 cytokines analyzed, 8 were found at higher levels (Table I) and 8 at lower levels (Table I) than the control cells (Fig. 6E), with a threshold of 2 fold change. Among the cytokines positively regulated during infection (Table I), some were involved in cartilage remodeling (chitinase 3, CD147 or EMMPRIN) or apoptosis (CD30, and IGFBP-3). Since CD147 is known to activate Matrix Metalloproteases (MMPs), we performed an RNA quantification of two MMPs involved in cartilage catabolism (MMP-3 and MMP-9).

**Table I:** Effect of CHIKV infection of pHCs in cytokine production. Supernatants from CHIKV-infected and uninfected HCs were analysed by a broad-spectrum enzyme-linked immunosorbent assay (Proteome Profiler Human XLCytokine array kit, R&D Systems), 6 hpi.

|  | Cytokine | Fold change (compared to mock infected) |
| --- | --- | --- |
| Upregulated | Serpin E1 (PAI1) | 13,8 |
|  | Chitinase-3 like 1 | 9 |
|  | IGFBP-3 | 4,6 |
|  | Complement factor D | 3,2 |
|  | FGF-19 | 3 |
|  | CD30 | 2,3 |
|  | IL-6 | 2,1 |
|  | EMMPRIN (CD147 or Basigin) | 2,1 |
| Downregulated | TGFα | -2,2 |
| | TNFα | -2,2 |
|  | TfR | -2,3 |
|  | VEGF | -2,6 |
|  | uPAR | -3 |
|  | IL-8 | -6,2 |
|  | MCP-1 | -6,7 |
|  | Thrombospondin-1 | -9,6 |

Both MMP’s RNA levels were higher in infected cells compared to control (Fig. 7A). As IGFBP-3 has been reported to induce chondrocyte apoptosis, we also investigated the induction of cell death by apoptosis following infection. The cell morphology and viability of infected pHCs were monitored by transmitted light microscopy and showed that infected cells underwent significant morphological changes and destruction of the cell layer increased massively between 24 and 72 hours.

**Figure 7.**
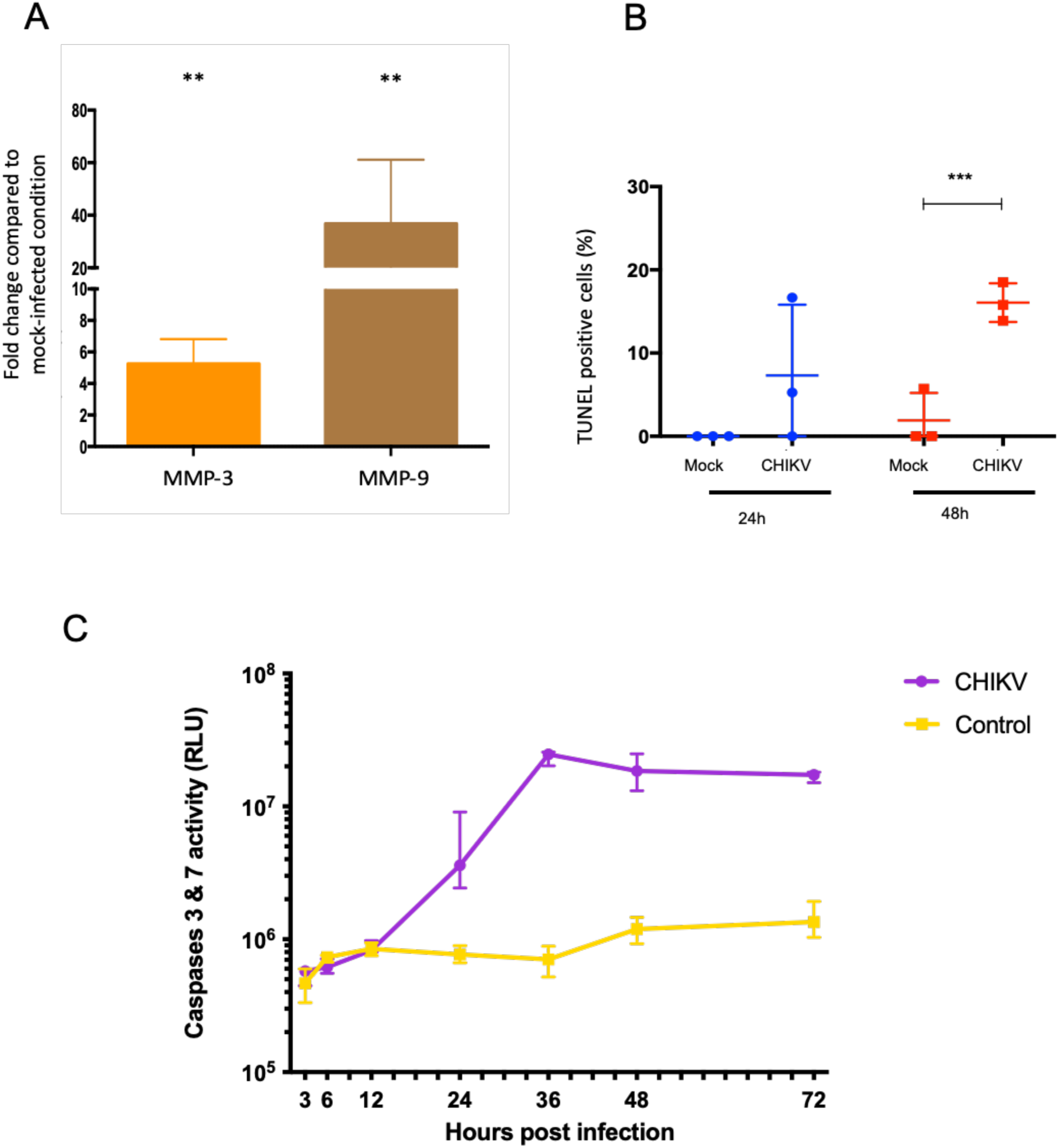
Viral induced alterations in HC biology. (A) Increased MMP-3 and MMP-9 mRNA synthesis after CHIKV infection of HCs. HCs were infected with CHIKV (MOI 1). At 48 hpi, MMP-3 and MMP-9 mRNAs were assessed by RT-qPCR. The amount of mRNA is expressed relative to uninfected controls, after normalization against a housekeeping gene (GAPDH). Histograms show the median value and interquartile range obtained in three independent experiments performed in triplicate. ns: not significant, **: p-value < 0.01 (non-parametric Mann-Withney test). (B) TUNEL labelling was performed and TUNEL positive cells were visualized using a fluorescence microscope. Average number of TUNEL-positive cells over three randomly observed fields (as a percentage of total) ns: not significant, ***: p-value < 0.001 (non-parametric Mann-Withney test). (C) Caspases 3 and 7 activity following CHIKV infection of pHCs. (MOI = 1). At different times pi, the cell monolayer was lysed and caspase 3 and 7 activity was measured (Caspase Glo 3/7 Assay Kit, Promega); bioluminescent signal is expressed in Units of Relative Luminescence (URL). The curves show the median value and interquartile range obtained in two independent experiments performed in triplicate. ns: not significant, **: p-value < 0.01 (non-parametric Mann-Withney test).

*In situ* cell death was confirmed by labelling with the TUNEL technique. At 24 hpi, we observed a slight increase in the number of TUNEL positive cells, although this was not significantly higher than control cells (Fig. 7B). At 48 hpi, 16.1% of the cells show positive TUNEL labelling, which was increased compared to the 1.9% basal level observed in control cells (Fig. 7B).

To further investigate the cell death mechanism involved, we measured the caspase 3/7 activity following pHCs infection with CHIKV and observed an important activation of the caspases starting at 24 hpi and reaching a plateau at 36 hpi (Fig. 7C).

## Discussion

The initial stage of our work involved adapting the adult murine model developed in 2010 by Gardner *et al*. [15] to our own experimental conditions using a new recombinant virus expressing NLuc recently developed in our laboratory. Indeed the bioluminescence live imaging approach has been shown valuable for studies of experimental CHIKV infection [21] and other arthritogenic alphaviruses such as Ross River virus [31]. We showed here the stability of the new recombinant virus over several passages on cells and observed a good correlation between bioluminescent signal and infectious virus titer. Since the reporter NLuc is expressed during the viral replication, the observation of a bioluminescent signal indicates active viral replication and allows here to overcome the limitation of viral RNA or antigens detection which can persist in tissues. Using the recombinant CHIKV-NLuc, we were able to observe a strong bioluminescent signal *in vivo* during the acute phase. Interestingly, the bioluminescent signal persisted beyond the acute stage, suggesting that complete replicating virus remained in the paws, inline with previous observations of Teo and al [21]. We observed bioluminescence reflecting viral replication in the joints of the inoculated paw (metatarsus, knee and hip). Over time, the signal was localized mainly in the metatarsi, similarly to the observations in humans showing a preferential impairment of the distal joints [34,35]. This could however also be a consequence of the choice of the inoculation site.

Moreover, we identified CHIKV infected tissues within the joint after isolation of cartilage from mice infected with the wild type virus. We were also able to isolate infected chondrocytes displaying dual immunoreactivity for the viral protein E and type II collagen. Immune cells infiltrating the joint or cartilage, synoviocytes or osteoblasts may have been present, but we were only able to identify infected chondrocytes within the cartilage tissue. Interestingly, the metatarsal joints appeared to be the main site for infected chondrocytes and this as late as 30 dpi. Other joints (e.g. knee) seemed to very rarely harbour infected cells, which could also be a consequence of the inoculation site, the metatarsi being the closest joints to the footpad. To our knowledge, our data demonstrate the first proof of *in viv*o chondrocytes infection with CHIKV during the acute phase and beyond. However, we believe it is crucial to confirm these results with other methods and/or different animal models. Indeed, our model was inoculated in the footpad with a high infectious dose, which does not reflect adequately natural infection.

In order to confirm our results in humans, and to be able to study more specifically the consequences of infection on the physiology of chondrocytes, we have subsequently used an *in vitro* model of primary chondrocytes isolated from samples taken from two healthy patients. We showed the ability of CHIKV to replicate efficiently in human chondrocytes leading to the production of new infectious viral particles. We also investigated the effect of infection on the chondrocyte’s physiology and observed significant changes in the secretion levels of several cytokines at 6 hpi. Sixteen cytokines were modulated, of which 8 were overexpressed. Interestingly 7 of the 8 overexpressed cytokines are found at high levels in patients with joint pathologies [36–42]. The role of these cytokines and their possible involvement in the pathogenesis of arthralgia and CHIKV infection was then investigated. E1 serpin, Chitinase 3-like 1, IGFBP-3, complement factor D, CD30, IL-6, and EMMPRIN emerge as highly expressed cytokines, potentially contributing to tissue degradation and immune response activation and indicating a disruption in cartilage metabolism regulation, favoring catabolism. Conversely, TNF-α, MCP-1, IL-8, VEGF, and thrombospondin were underexpressed, highlighting the intricate and complex cytokine interplay during CHIKV infection. We futher explored the effect of CHIKV infection on human chondrocytes and showed an activation of matrix metalloproteinases (MMP-3 and MMP-9), indicating the establishment of a mechanism promoting cartilage breakdown. Finally, using a TUNEL assay, we showed CHIKV induced apoptosis in chondrocytes, confirming the effect of factors like IGFBP-3 on the cells physiology after infection.

Interestingly, *in vitro* observations and especially the rapid cell death following chondrocytes infection do not align with *in vivo* outcomes, hinting at potential protective factors within the joint environment. One hypothesis would be related to the organization of the cartilage and its relative paucity in cells, that could limit viral replication, contrary to the replication observed *in vitro*. However, while *in vitro* observations provide insights, the complex *in vivo* scenario can also be attributed to paracrine regulations by joint cells, underscoring again the intricate dynamics within the joint environment.

In summary, we showed here the susceptibility of chondrocytes to CHIKV infection both *in vitro* and *in vivo*, during and past the acute phase. We observed modifications in the cytokines secretion and the activation of apoptosis, leading to a dysregulation in the cartilage metabolism. Our study sheds light on the intricate network of cytokines, MMP and apoptotic pathways affected by CHIKV infection in the joints, emphasizing the multifaceted impact on chondrocytes and cartilage.

