## Supplementary figure 1 for "A recombinant CHIKV-NLuc virus identifies chondrocytes as target of Chikungunya virus in a immunocompetent mouse model"

### Supplementary material

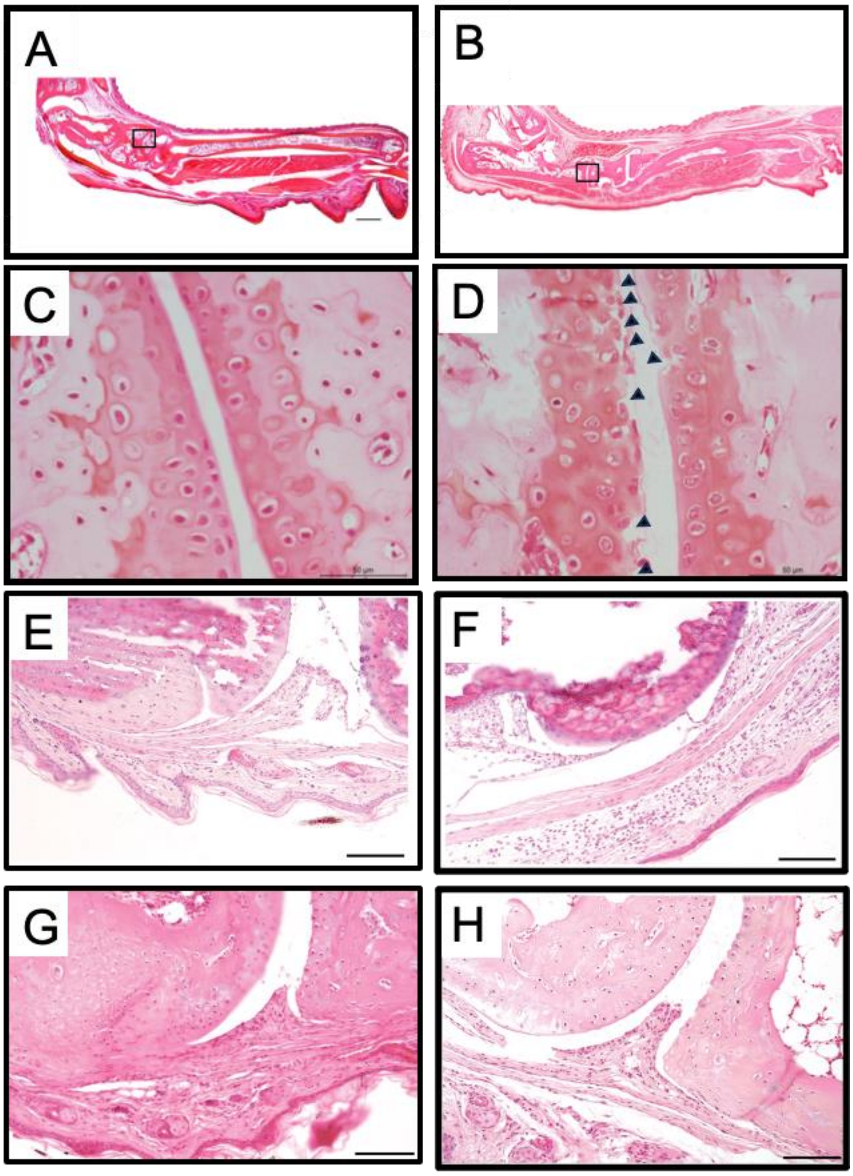


**Supplementary figure 1.** Joint damage after CHIKV infection in mice.
Mice were inoculated in the plantar pad of the right hind paw with CHIKV ($10^5$) or diluent alone (control). Six days after infection, mice were killed and the inoculated paws were harvested and fixed in formalin, then stained with safranin solution, allowing cartilage staining. Longitudinal sections of a mouse paw inoculated with diluent alone (A, C) or with CHIKV (B, D) are shown, together with details of the articular cartilage. Black arrows indicate erosive cartilage lesions (Scale Bar: 50 mm). When stained with hemalun/eosin solution, 6 days p.i., preserved structure of the joint was observed in the uninfected control (E); in contrast, joint lesions were visible in infected mice killed 6 days p.i., (F), 12 days p.i., (G) and 30 days p.i.. (Scale Bar: 100µm).
